# Lipid peroxidation drives liquid-liquid phase separation and disrupts raft protein partitioning in biological membranes

**DOI:** 10.1101/2023.09.12.557355

**Authors:** Muthuraj Balakrishnan, Anne K. Kenworthy

**Author notes:** McKetta Department of Chemical Engineering, University of Texas at Austin, Austin, Texas 78712, United States (Muthuraj Balakrishnan).

## Abstract

The peroxidation of membrane lipids by free radicals contributes to aging, numerous diseases, and ferroptosis, an iron-dependent form of cell death. Peroxidation changes the structure, conformation and physicochemical properties of lipids, leading to major membrane alterations including bilayer thinning, altered fluidity, and increased permeability. Whether and how lipid peroxidation impacts the lateral organization of proteins and lipids in biological membranes, however, remains poorly understood. Here, we employ cell-derived giant plasma membrane vesicles (GPMVs) as a model to investigate the impact of lipid peroxidation on ordered membrane domains, often termed membrane rafts. We show that lipid peroxidation induced by the Fenton reaction dramatically enhances phase separation propensity of GPMVs into co-existing liquid ordered (raft) and liquid disordered (non-raft) domains and increases the relative abundance of the disordered, non-raft phase. Peroxidation also leads to preferential accumulation of peroxidized lipids and 4-hydroxynonenal (4-HNE) adducts in the disordered phase, decreased lipid packing in both raft and non-raft domains, and translocation of multiple classes of proteins out of rafts. These findings indicate that peroxidation of plasma membrane lipids disturbs many aspects of membrane rafts, including their stability, abundance, packing, and protein and lipid composition. We propose that these disruptions contribute to the pathological consequences of lipid peroxidation during aging and disease, and thus serve as potential targets for therapeutic intervention.

## Introduction

The lipid composition of biomembranes is tuned to support the function of membrane proteins and cellular processes [1]. Genetic defects and environmental insults that disrupt the normal repertoire of lipids can have profound consequences [2]. One such insult is lipid peroxidation, a process driven by unregulated oxidative stress.

During lipid peroxidation, lipids containing carbon-carbon double bonds, particularly lipids containing polyunsaturated fatty acids (PUFAs) which have many such double bonds, undergo free radical attack by oxygen radicals [3–6]. Lipid peroxidation is mediated via reactive oxygen species (ROS) and can proceed by both non-enzymatic (e.g. the Fenton reaction) and enzymatic mechanisms (e.g. lipoxygenases/oxidases). In the Fenton reaction, hydrogen peroxide (H_2_O_2_) reacts with redox metal, such as iron, to generate a hydroxyl radical [5, 7, 8]. The hydroxyl radical reacts with almost all biological molecules in living cells, forming products which can themselves further damage biological structures. PUFA-containing lipids are particularly susceptible to this chain-reaction because of their numerous carbon double bonds [3–6].

Lipid peroxidation produces two main types of products [3–6]. The primary products are hydroperoxide lipids. Their hydroperoxidized acyl chains becomes more hydrophilic and tend to associate with the lipid-water interface [9–11]. Their presence in membranes ultimately increases membrane permeability and impacts other membranes properties such as packing, fluidity, and viscosity [12–20]. Further oxidation generates secondary products such as reactive aldehydes, ketones, alcohols and ethers. The best studied secondary products of lipid peroxidation include the bioactive molecules 4-hydroxynonenal (4-HNE), a product of arachidonic acid, and 4-hydroxyhexenal (4-HHE), generated from docosahexaenoic acid. These reactive compounds form molecular adducts with lipids, proteins, DNA and other biomolecules, thereby disrupting their normal functions [21–23].

The biological consequences of lipid peroxidation are substantial. Lipid peroxidation triggers ferroptosis, an iron-dependent form of cell death [24–26]. It also contributes to aging, neurodegenerative diseases such as Alzheimer’s and Parkinson’s disease, and cancer [27–31]. To combat the deleterious effects of lipid peroxidation, efforts are underway to develop and optimize inhibitors of the process [32–34]. Conversely, lipid peroxidation can be harnessed and exploited as a tool in photodynamic therapy to treat cancer and other diseases [35, 36]. Given these important roles of lipid peroxidation in biology, there is a substantial need to not only understand the biochemical mechanisms that underlie the process, but also to uncover how lipid peroxidation impacts the structure and function of cells.

Phase separation plays an important role in the function of biological systems [37–41]. For many years, the plasma membrane of cells has been proposed to be capable of undergoing a form of liquid-liquid phase separation similar to that observed in ternary lipid mixtures consisting of a saturated lipid with a high melting temperature, an unsaturated lipid with a low melting temperature, and cholesterol, resulting in the formation of co-existing liquid ordered (raft) and liquid disordered (non-raft) domains [39, 42–44]. In cells, raft domains are transient and dynamic, enriched in sphingolipids, cholesterol, and lipids with saturated acyl chains, and contain a subset of membrane proteins, whereas non-raft domains are enriched in unsaturated and polyunsaturated lipids as well as membrane proteins that prefer to reside in a disordered environment [45–49]. Rafts have been implicated in a wide range of cellular functions such as cell signaling, membrane trafficking, cellular adhesion and motility, and other essential cellular functions [45, 46, 48]. There is thus considerable interest in understanding their contributions to health and disease [46, 50–55].

Interestingly, in model systems the presence of oxidized lipids has been reported to impact lipid phase separation, in some cases promoting raft formation while disrupting rafts in others [56–60]. Furthermore, polyunsaturated lipids, the main target of lipid peroxidation, are known to regulate raft formation and composition [49, 61–70]. Yet, how lipid peroxidation impacts the formation, properties, and function of rafts in biological membranes has yet to be investigated. Here, we show that lipid peroxidation has profound consequences on the stability, abundance, packing, and protein and lipid composition of both raft-like and non-raft domains in biological membranes.

## Results

### GPMVs are a useful model to study lipid peroxidation

Current models suggest the lipid composition of the plasma membrane of mammalian cells is tuned to position the lipids close to a miscibility critical point, which enables them to undergo phase separation to form co-existing ordered (raft) and disordered (non-raft) domains in response to small perturbations [44, 48, 71]. Under physiological conditions, these domains are nanoscopic in size [46]. This makes them difficult to detect in living plasma membranes. We therefore turned to giant plasma membrane vesicles (GPMVs) as a model to study the effects of lipid peroxidation on rafts. Unlike in the plasma membrane of live cells, in cell-derived GPMVs, co-existing raft and non-raft domains are large enough to be directly visualized by fluorescence microscopy [72–76]. Formation of these two macroscopic phases occurs below the miscibility transition temperature and is thought to reflect the presence of much smaller domains at physiological temperatures [71, 77, 78].

GPMVs are generated by treating mammalian cells with combinations of blebbing reagents, leading to their release from the cell surface. They can then be harvested, subjected to various treatments, and incubated with fluorescent dyes to selectively label either raft or non-raft domains or probe their biophysical properties [75, 76]. For our studies, GPMVs were isolated from HeLa cells. At room temperature, a major fraction of HeLa cell derived GPMVs typically exhibit phase separation [79–83]. This makes them a useful system to investigate factors that influence membranes liquid-liquid phase separation and the affinity of proteins and lipids for raft-like versus non-raft membrane domains [72–76, 84, 85]..

To induce lipid peroxidation in GPMVs, we applied a chemical approach based on the Fenton reaction. In this reaction, cations of a transition metal such as iron interact with hydrogen peroxide to form a hydroxyl radical [8, 86]. Here, we utilized Fe(II) as an electron source and cumene hydroperoxide as a stable, lipophilic oxidizing agent [87–89] (**Fig. 1A**). In this reaction, the reduced form of Fe(II) is oxidized to Fe(III) by the Fenton reaction with cumene hydroperoxide to form lipophilic cumoxyl radicals. Cumoxyl radicals can also react with another cumene hydroperoxide molecules to yield cumoperoxyl radicals, which in turn leads to lipid peroxidation (reviewed in [23]). We chose to use a 1:10 metal: cumene hydroperoxide ratio based on a previous study [90].

**Figure 1.**
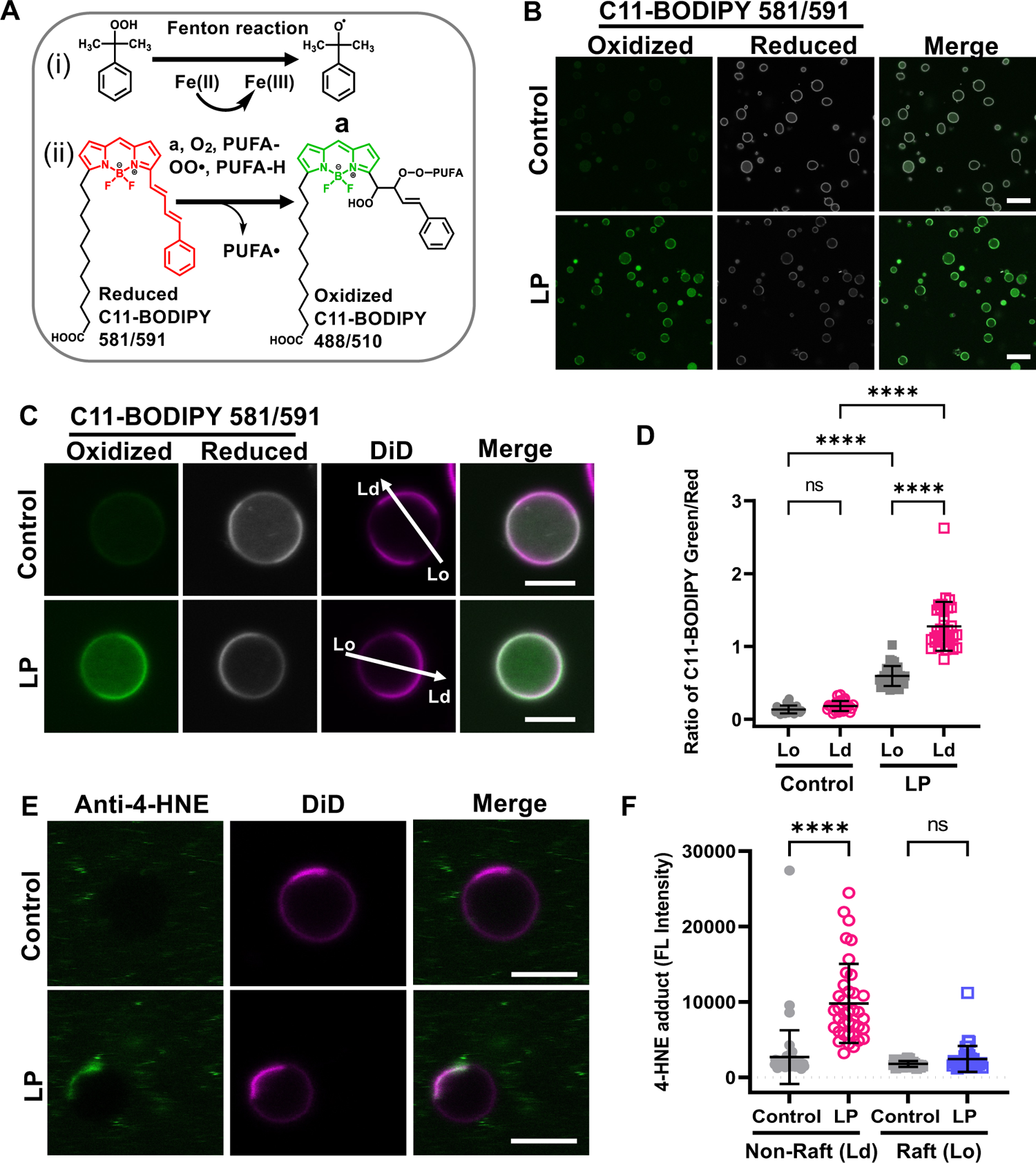
Induction and detection of lipid peroxidation in GPMVs. (**A)** (i) Cumene hydroperoxide in the presence of transition metal iron Fe(II) produces cumoxyl radicals via the Fenton reaction (a). (ii) Cumoxyl radicals (a) abstract a hydrogen (H) from a polyunsaturated lipid (PUFA-H) generating a lipid radical (PUFA•) that reacts immediately with oxygen, generating PUFA peroxy radicals (PUFA-OO•). The PUFA peroxy radicals react with C11-BODIPY 581/591, causing deconjugation of C11-BODIPY and a blue shift of the emission wavelength to 510 nm. **(B)** GPMVs were left untreated or incubated with 50 μM Fe(II) and 500 μM cumene hydroperoxide to induce lipid peroxidation, labeled with 1 μM of BODIPY 581/591, allowed to settle for 15-20 min, and imaged at RT using confocal microscopy. Scale bars, 20 μm. (**C**) Representative images of individual GPMVs labeled with C11-BODIPY 581/591 under control conditions or following lipid peroxidation. GPMVs were co-labeled with DiD to mark the position of the disordered domains. The arrows show examples of the position of lines used to analyze the fluorescence intensity in Lo and Ld domains. Scale bars, 5 μm. (**D**) Ratio of green (oxidized): red (reduced) BODIPY 581/591 fluorescence in ordered versus disordered domains under control conditions and following lipid peroxidation. Each data point corresponds to an individual GPMV. Error bars show mean ± SD for 27-33 GPMVs. **(E)** GPMVs were either left untreated or subjected to lipid peroxidation, immunolabeled with an anti-4-HNE antibody and Alexa-488 secondary antibody, and then stained using DiD. Examples of representative GPMVs are shown. Scale bars, 20 μm. **(F)** Quantification of immunostaining of 4-HNE levels. Fluorescence intensity is reported in arbitrary units. Each data point corresponds to an individual GPMV. Error bars show mean ± SD. P values were determined by unpaired one-way ANOVA with Sidak’s multiple comparison test, Alpha = 0.05 (95% confidence level), ****, P<0.0001; n.s., not significant. Data in D and E were pooled across 2 independent experiments.

To detect lipid peroxidation, we used the lipid peroxidation sensor C11-BODIPY581/591 (C11-BODIPY) [89, 91]. C11-BODIPY binds membranes and exhibits a fluorescence emission peak shift from 590 nm in its reduced state (red) to 510 nm in its oxidized state (green) induced by lipid peroxidation (**Fig. 1A**). In control GPMVs labeled with C11-BODIPY, primarily red fluorescence was observed (**Fig. 1B** and **Fig S1A**). In contrast, after incubating GPMVs with Fe(II) (50 μM) and cumene hydroperoxide (500 μM), the fluorescence signal from the reduced form of C11-BODIPY (red) decreased, and signal from the oxidized form of C11-BODIPY (green) concomitantly increased (**Fig. 1B**). No fluorescence was observed in GPMVs incubated with lipid peroxidation reagents in the absence C11-BODIPY (**Fig. S1**). Furthermore, when either Fe(II) or cumene hydroperoxide were omitted from the reaction, C11-BODIPY remained in a reduced state, indicating both Fe(II) and cumene hydroperoxide are required to drive lipid peroxidation (**Fig. S1**). These results suggest GPMVs can be used to investigate the effects of lipid peroxidation on cell plasma membranes.

In GPMVs subjected to lipid peroxidation, C11-BODIPY fluorescence tended to be enriched in one domain in phase separated GPMVs (**Fig. 1C**). To determine whether this partitioning corresponds to the ordered or disordered domain, GPMVs were labeled with both C11-BODIPY, and the disordered phase marker DiD [92, 93] prior to imaging. In both control GPMVs and GPMVs that had undergone lipid peroxidation, the phase containing reduced C11-BODIPY was depleted in DiD fluorescence, suggesting it corresponds to the ordered phase. Areas with high levels of green fluorescence (oxidized C11-BODIPY), on the other hand, typically also contained more intense DiD fluorescence, implying they correspond to disordered domains (**Fig. 1C and S2**). Quantification of the ratio of green (oxidized): red (reduced) C11-BODIPY fluorescence in Lo and Ld domains in populations of GPMVs revealed significantly higher ratio of green versus red C11-BODIPY fluorescence in GPMVs subjected to lipid peroxidation relative to controls (**Fig. 1D**). Interestingly, the green to red ratio was significantly higher in the disordered domains in treated GPMVs (**Fig. 1D**). This suggests that levels of peroxidized lipids are higher in disordered than ordered domains.

Lipid peroxidation generates a variety of aldehyde products that can covalently modify proteins and lipids [21, 22]. To test whether these products were being formed under the conditions of our experiments, we examined the distribution of aldehyde 4-hydroxynonenal (4-HNE), one of the best studied bioactive products of lipid peroxidation [23, 30]. Immunostaining of GPMVs showed that 4-HNE products were present after, but not before, lipid peroxidation (**Fig. 1E, F**; **Fig. S3**). Notably, 4-HNE staining colocalized with the disordered phase marker DiD, similar to the distribution of peroxidized lipids (**Fig. 1E, F**).

Together, these results suggest that the Fenton reaction drives lipid peroxidation in GPMVs as reported by BODIPY-C11 oxidation and the accumulation of 4-HNE, and that these products preferentially accumulate in Ld domains.

### Lipid peroxidation enhances phase separation in GPMVs

The propensity of GPMVs to phase separate into coexisting raft and non-raft domains depends on membrane lipid composition as well as the physicochemical properties of raft and non-raft assemblies, such as the order of each phase [68, 74]. We thus next assessed the effects of lipid peroxidation on phase separation. GPMVs were labeled with NBD-DSPE (ordered domain marker) and DiD (disordered domain marker) to visualize ordered and disordered domains simultaneously [72, 82]. In control experiments, approximately 40% of vesicles exhibit phase separation (**Fig. 2A, B**). Remarkably, after lipid peroxidation, almost all vesicles (>90%) were phase-separated (**Fig. 2A, B**). Similar results were obtained using GPMVs generated from RPE1 cells (**Fig. S3**), suggesting lipid peroxidation enhances phase separation in a cell type-independent manner. For consistency, unless otherwise indicated, in all subsequent experiments we utilized HeLa cell-derived GPMVs.

**Figure 2.**
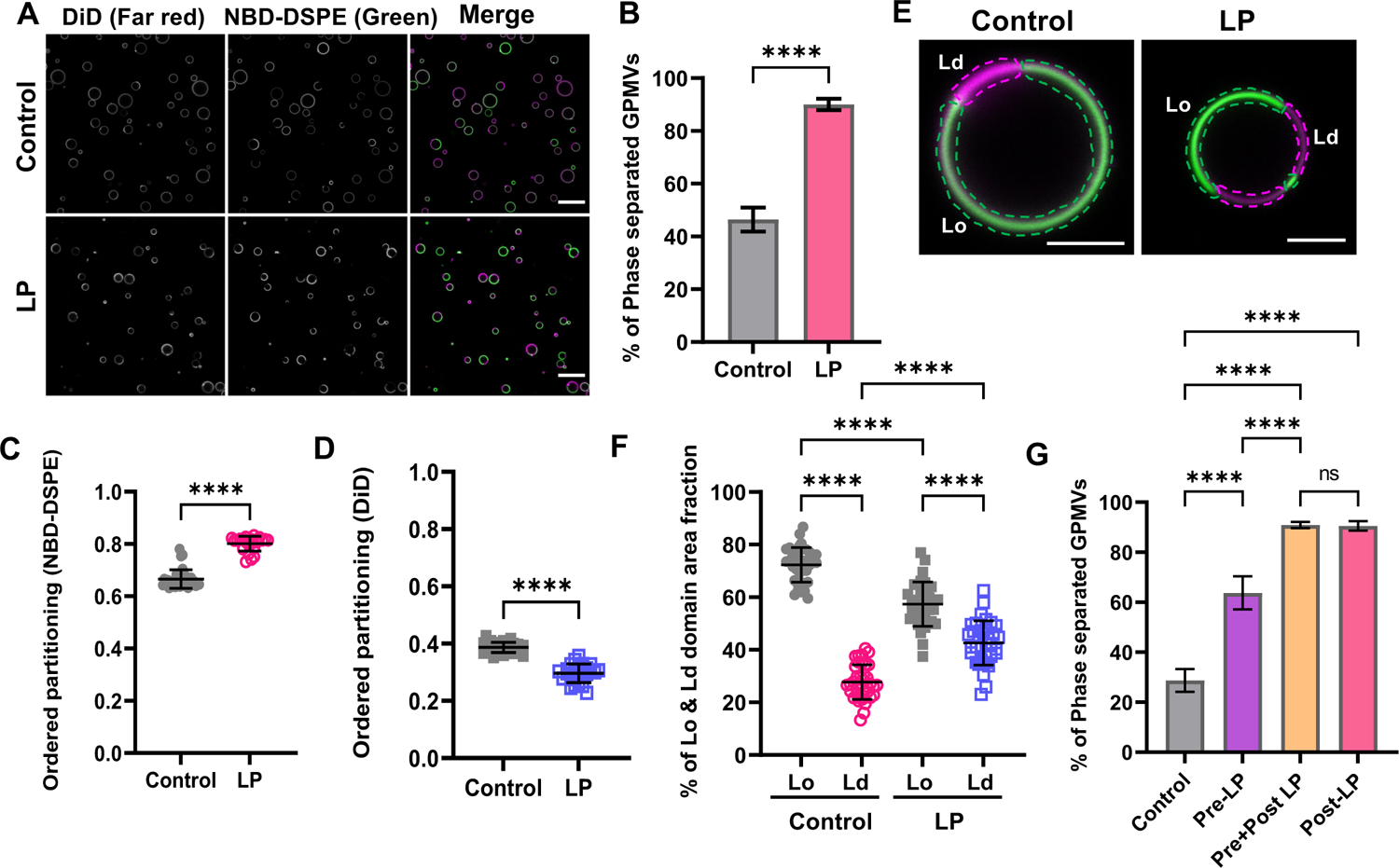
Lipid peroxidation increases the percentage of phase separated vesicles and the area fraction of disordered domains. (**A**) GPMVs were either left untreated (control) or subjected to lipid peroxidation (LP). They were then labeled sequentially with NBD-DSPE (green) and DiD (magenta) prior to imaging at RT using confocal microscopy. Scale bar, 20 μm. (**B**) Quantification of the percentage of phase separated GPMVs for control versus lipid peroxidation conditions. The % of phase separated GPMVs was calculated using the green channel using VesA software. Data are presented as mean ± SD for >1000 GPMVs per group. Data were pooled across 8 independent experiments. (**C, D**) Impact of lipid peroxidation on ordered partitioning of raft (NBD-DSPE) and non-raft (DiD) reporter dyes. Points in C and D represent >100 GPMVs in each group. ****, P < 0.0001 using unpaired two-tailed t-test. Data are representative of 8 independent experiments. Each datapoint is for an individual field of GPMVs containing >50 GPMVs. (**E**) Illustration of how the area fraction of raft (green) and non-raft (magenta) domains was quantified for representative GPMVs. Scale bars, 5 μm. (**F**) Effect of lipid peroxidation on the area fraction of Lo and Ld domains. Each data point corresponds to an individual GPMV. Bars show the mean ± SD for two independent experiments. P values were determined by unpaired one-way ANOVA with Sidak’s multiple comparison test, Alpha = 0.05 (95% confidence level). ****, P<0.0001. (**G**) HeLa cells were left untreated (control) or pretreated with lipid peroxidation reagents for 30 min at RT (pre-LP) prior to GPMV preparation. For comparison, a population of control GPMVs and pre-LP GPMVs were subsequently incubated with lipid peroxidation reagents (post-LP and pre + post LP, respectively). All GPMVs were then labeled with NBD-DSPE and DiD and imaged using confocal microscopy. The % of phase separated GPMVs was calculated using the green channel using VesA software. Data are presented as mean ± SD. P values were determined by unpaired one-way ANOVA with Sidak’s multiple comparison test, Alpha = 0.05 (95% confidence level) ****, P<0.0001; n.s., not significant. Data are representative of 3 independent experiments for >100 GPMVs per group.

Changes in the properties of the raft or non-raft domains are also predicted to alter the partitioning of lipid probes [82]. We thus next quantified the effect of lipid peroxidation on the preference of the fluorescent domain markers for raft versus non-raft domains, defined as P_ordered_ [81, 83],

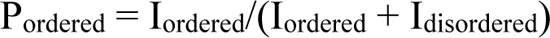

where I_ordered_ is the fluorescence intensity of the molecule of interest in the Lo domain and I_disordered_ is the fluorescence intensity in the Ld-like domain, with the underlying assumption that the fluorescence intensity is proportional to the concentration of fluorescent molecules in the two environments (**Fig. S4**). P_ordered_ values range from 0 to 1, where P_ordered_ > 0.5 indicates that the marker prefers the ordered phase and P_ordered_ < 0.5 means that the domain marker prefers the disordered phase. We found that P_ordered_ increased for NBD-DSPE in response to lipid peroxidation, whereas P_ordered_ for DiD decreased (**Fig. 2C, D**). These findings illustrate changes in the properties and compositions of the domains in response to lipid peroxidation.

We next asked whether lipid peroxidation impacts the relative abundance of raft versus non-raft domains. We measured the area fractions of Lo and Ld domains across multiple GPMVs and experiments (**Fig. 2E, F**). Lipid peroxidation decreased the area fraction of Lo domains from ∼0.7 to ∼0.6 (**Fig. 2E, F**) while the fraction of Ld domains significantly increased from ∼0.3 to ∼0.4 (**Fig. 3E, F**). This suggests that either there is a shift in the total number of lipids present in the raft phase versus the non-raft phase or that the average area per molecule increases more strongly in the disordered than the ordered phase in response to lipid peroxidation.

**Figure 3.**
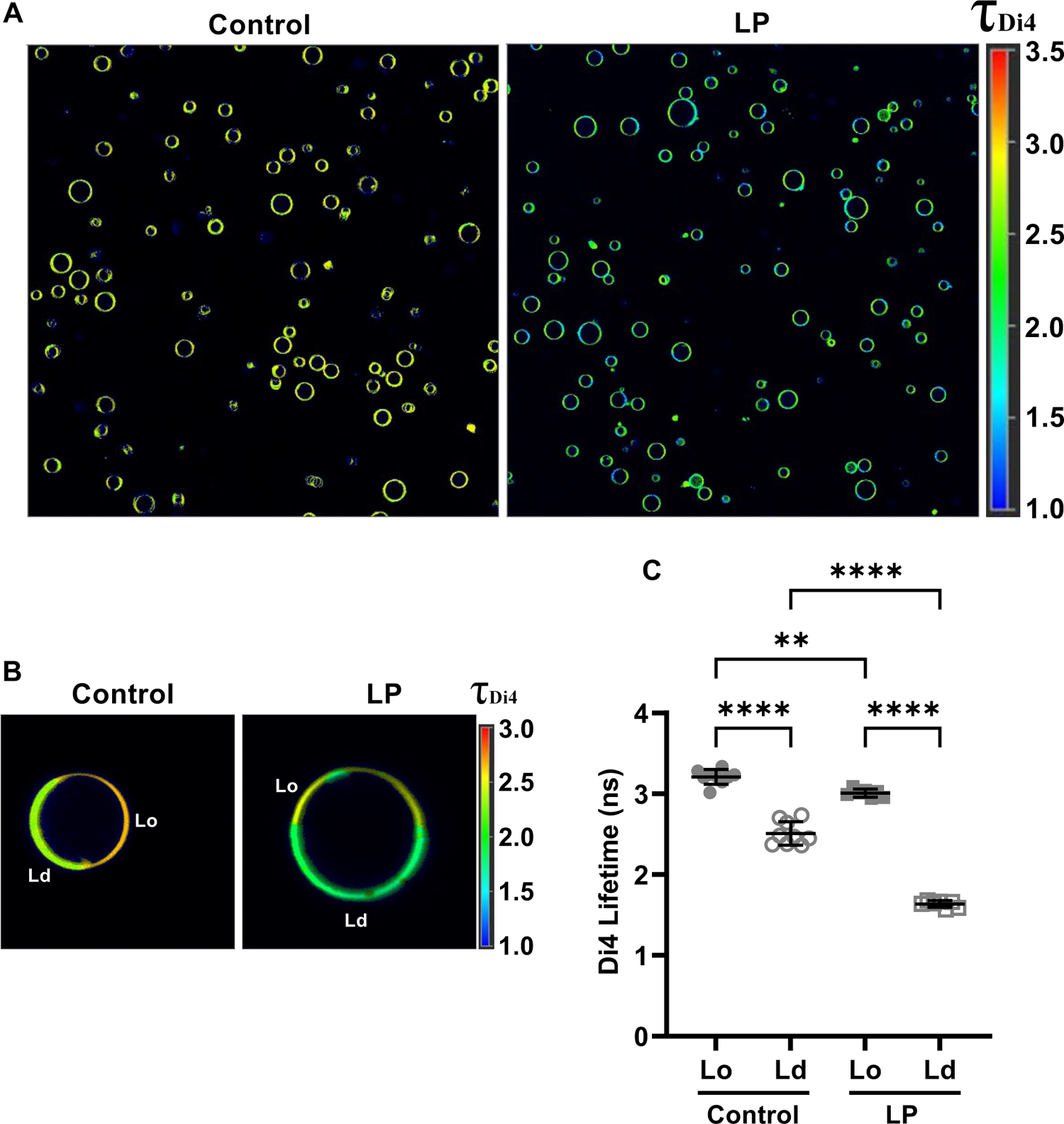
Lipid peroxidation decreases lipid packing in both ordered and disordered domains. (**A**) Fluorescence lifetime imaging microscopy (FLIM) images of GPMVs labeled with Di4 under control conditions and following lipid peroxidation. Lookup table shows Di4 lifetime (ns). (**B**) Representative FLIM images of individual GPMVs, highlighting differences in Di4 lifetime in Lo and Ld domains. Lookup table shows Di4 lifetime (ns). (**C**) Quantification of Di4 lifetimes in Ld and Lo domains for control GPMVs and GPMVs subjected to lipid peroxidation. For the control sample, individual data points correspond to mean values for a field of >10-15 GPMVs taken from a single lifetime image; total number of control GPMVs = 104. For the LP sample, individual data points correspond to mean values for a field of >50 GPMVs taken from a single lifetime image; total number of LP GPMVs = 433. Error bars show mean ± SD. P values were determined by unpaired one-way ANOVA with Sidak’s multiple comparison test, Alpha = 0.05 (95% confidence level), ****P<0.0001; n.s., not significant. Data are representative of two independent experiments.

The above experiments were performed by directly subjecting GPMVs to conditions that induce lipid peroxidation. We next tested whether similar effects were observed in GPMVs derived from cells subjected to oxidative stress. For these experiments, we treated HeLa cells with lipid peroxidation reagents for 30 min prior to GPMV preparation. GPMVs again showed evidence for perturbed membrane phase behavior, including a significantly increased percentage of phase separated GPMVs (**Fig. 2G**). The percentage of phase separation was increased further by post-treating these GPMVs with lipid peroxidation reagents (**Figure 2G**). For simplicity, further experiments were carried out with GPMVs subjected directly to lipid peroxidation.

Taken together, these results suggest that lipid peroxidation has a profound effect on plasma membrane phase behavior, dramatically enhancing phase separation, altering the proportion of ordered and disordered domains, and modulating the partitioning of fluorescent lipids. To gain further insights into the effects of lipid peroxidation on the membrane, we next examined lipid packing.

### Lipid peroxidation decreases lipid packing in both ordered and disordered domains

Lipid peroxidation generates products such as truncated lipids that perturb lipid packing in synthetic model membranes [10, 11, 14–16, 19, 20]. To test whether similar effects are relevant in biological membranes, we turned to environment-sensitive fluorescent probes that sense lipid packing. Specifically, we monitored the lifetime of the fluorescent reporter Di4 (Di-4-ANEPPDHQ) [94, 95], which has been widely shown to be dependent on membrane lipid packing, with lower Di4 lifetimes indicative of looser packing and more fluid membranes, and vice versa [96].

Representative fluorescence lifetime imaging microscopy (FLIM) images of control and treated GPMVs are shown in Figure 3A. Based on the literature [96], domains with the higher lifetime correspond to the Lo domain and domains with the lower lifetime represent the Ld domain. Di4 lifetime decreased substantially in both domains in response to lipid peroxidation (**Fig. 3A, B**). In Lo domains, Di4 lifetime decreased from 3.26 ± 0.05 ns to 3.02 ± 0.07 ns following lipid peroxidation (**Fig. 3C**). Lipid peroxidation also decreased Di4 lifetime in the disordered domains from 2.46 ± 0.09 ns to 1.62 ± 0.08 ns (**Fig. 3C**). Notably, a larger change in Di4 lifetime was observed in Ld domains (Δτ = 0.84 ns) than in Lo domain (Δτ = 0.24 ns) following lipid peroxidation, indicative of more dramatic disruption of lipid packing in Ld domains. This is likely due to the accumulation of higher levels of peroxidized lipids in Ld domains (**Fig. 1**).

### Raft-preferring proteins translocate to non-raft domains in response to lipid peroxidation

A major function of rafts in biological membranes is to laterally compartmentalize proteins [46]. To facilitate sorting into or out of rafts, the structural features of proteins are tuned to match the fundamental properties of the membrane bilayer of each phase, including its thickness and lipid packing [47, 97, 98]. Polyunsaturated lipids have been proposed to regulate the affinity of proteins for ordered domains [62, 65, 70, 99–102]. However, the effect of peroxidation on protein sorting into rafts is largely unknown. We thus set out to test whether the changes we observed in the properties of the raft and non-raft phases could change protein partitioning.

We first studied a GPI-anchored protein, YFP-GL-GPI [83, 85]. GPI-anchored proteins are normally targeted to rafts by the saturated acyl chains of the GPI-anchor and are used extensively as markers for raft domains [46, 103]. As expected, YFP-GL-GPI partitioned to the ordered raft phase in untreated GPMVs (**Fig. 4**). Lipid peroxidation dramatically disrupted this partitioning, leading to YFP-GL-GPI accumulation instead in the non-raft membrane regions (**Fig. 4A**). Similar results were obtained using several non-raft reporter dyes, confirming this shift was not due to changes in the localization of the marker dyes themselves (**Figure S6A, B**). The expulsion of YFP-GL-GPI from rafts was quantified as a dramatic decrease in P_ordered_ in response to lipid peroxidation (**Fig. 4B**). Thus, peroxidation-induced changes disrupt proper partitioning of the GPI-anchor to ordered membrane domains.

**Figure 4.**
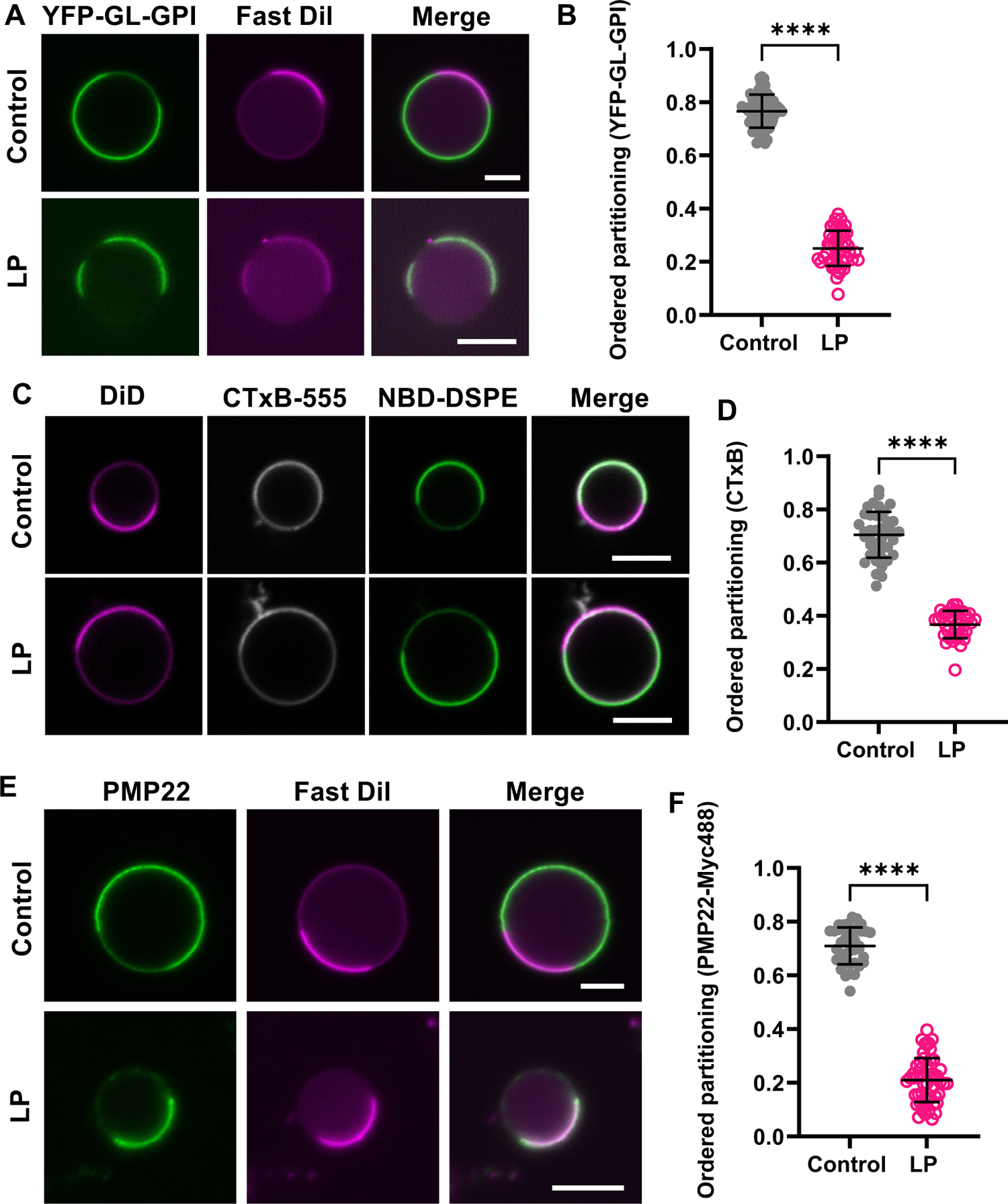
Raft-preferring proteins redistribute to non-raft domains in response to lipid peroxidation. (**A**) Representative images of YFP-GL-GPI in HeLa cell GPMVs. GPMVs were stained with Fast DiI after lipid peroxidation. (**B**) Impact of lipid peroxidation on ordered partitioning of YFP-GL-GPI. Each data point corresponds to an individual GPMVs. Data are presented as mean ± SD for n =47-68 GPMVs. **(C)** Representative images of CTxB-Alexa 555 labeling of COS-7 cell-derived GPMVs. After lipid peroxidation, GPMVs were sequentially labeled with CTxB-Alexa 555, NBD-DSPE, and DiD. (**D**) Impact of lipid peroxidation on ordered partitioning of CTxB-Alexa 555. Data are presented as mean ± SD for 37-41 GPMVs. (**E**) Representative images of PMP22 in HeLa cell GPMVs. GPMVs were labeled with an Alexa488-labeled anti-myc antibody, subjected to lipid peroxidation, and then labeled with Fast DiI. (**F**) Impact of lipid peroxidation on ordered partitioning of PMP22. Data are presented as mean ± SD for 39-56 GPMVs. **, P<0.01; ****P < 0.0001 for unpaired two-tailed t-test. All data are representative of 2 independent experiments. Scale bars, 5 μm.

To test the generality of these findings, we examined two other raft-associated proteins, cholera toxin B-subunit (CTxB) and Peripheral Myelin Protein 22 (PMP22). CTxB is a well-known raft marker used to study the functions and properties of rafts domains [104]. It binds up to five molecules of its glycolipid receptor, ganglioside GM1 [104]. Naturally occurring forms of GM1 are typically enriched in rafts [105], and binding of CTxB to GM1 further enhances its association with raft domains via a clustering mechanism [106]. In contrast, PMP22 is a multipass transmembrane protein [107]. PMP22 also preferentially partitions in ordered domains in the plasma membrane, by mechanisms that are not yet clear [81]. For our experiments, we utilized an N-glycosylation mutant of PMP22, N41Q-PMP22, that is targeted to the plasma membrane more efficiently that the wild type form of the protein [81]. N41Q-PMP22 (referred to hereafter as PMP22 for simplicity) contains a c-myc tag inserted into the second extracellular loop and can be visualized by labeling GPMVs with fluorescently labeled myc antibodies [81].

To study the impact of lipid peroxidation on the phase preference of CTxB bound to endogenous GM1, we generated GPMVs from COS-7 cells, which bind high levels of CTxB [106, 108]. As expected, CTxB colocalized with the raft marker NBD-DSPE in untreated GPMVs (**Fig. 4C**). In contrast, when CTxB was added to GPMVs subjected to lipid peroxidation, it localized predominantly to non-raft domains, corresponding to a significant decrease in P_ordered_ (**Fig. 4C, D**). Similar results were obtained for PMP22 in HeLa cell-derived GPMVs (**Fig. 4E, 4F**).

Together, these results demonstrate that lipid peroxidation causes multiple classes of raft-preferring proteins, including a GPI-anchored protein, glycolipid-binding protein, and multi-pass transmembrane protein to shift out of the more ordered phase into the more disordered membrane environment. Thus, lipid peroxidation alters not only the lipid composition of both raft and non-raft domains, but also their protein composition due to the mis-localization of raft proteins.

### Non-raft proteins remain associated with the disordered phase following lipid peroxidation

We next examined the impact of lipid peroxidation on proteins that preferentially reside in non-raft domains. Most examples of non-raft proteins are transmembrane proteins. As a test case, we studied the amyloid precursor protein (APP) and its cleavage product C99, key players in an amyloidogenic pathway linked to Alzheimer’s disease [109]. In the amyloidogenic pathway, APP is cleaved by β-secretase to yield C99, which is subsequently processed by γ-secretase to generate amyloidogenic Aβ peptides [110, 111]. Although these processing events have long been thought to occur in raft-like environments [112–114], we recently found that C99-EGFP preferentially localizes in disordered regions of the plasma membrane [83]. Lipid peroxidation has also been implicated in the progression of Alzheimer’s disease [31, 115–117]. We therefore wondered if lipid peroxidation would enhance the partitioning of APP or C99 into rafts.

To test this, we examined the phase preference of GFP-tagged APP and C99 in GPMVs. To prevent APP and C99 from being cleaved by γ-secretase, we included the γ-secretase inhibitor DAPT throughout the experiment [83]. Under control conditions, APP and C99 both colocalized with the DiD-enriched phase, corresponding to the more disordered region of the membrane (**Fig. 5A, C**). Interestingly, both APP and C99-EGFP remained exclusively associated with disordered regions of the membrane following lipid peroxidation (**Fig. 5A-D**). To investigate whether this is a general characteristic of non-raft proteins, we studied a GFP-tagged form of transferrin receptor, TfR-GFP [118]. TfR-GFP localized primarily in disordered domains under control conditions as expected (**Fig. 5E**) and remained enriched in non-raft regions of the membrane following lipid peroxidation (**Fig. 5E, F**). Together, these findings suggest that non-raft proteins continue to partition in this environment in peroxidized membranes.

**Figure 5.**
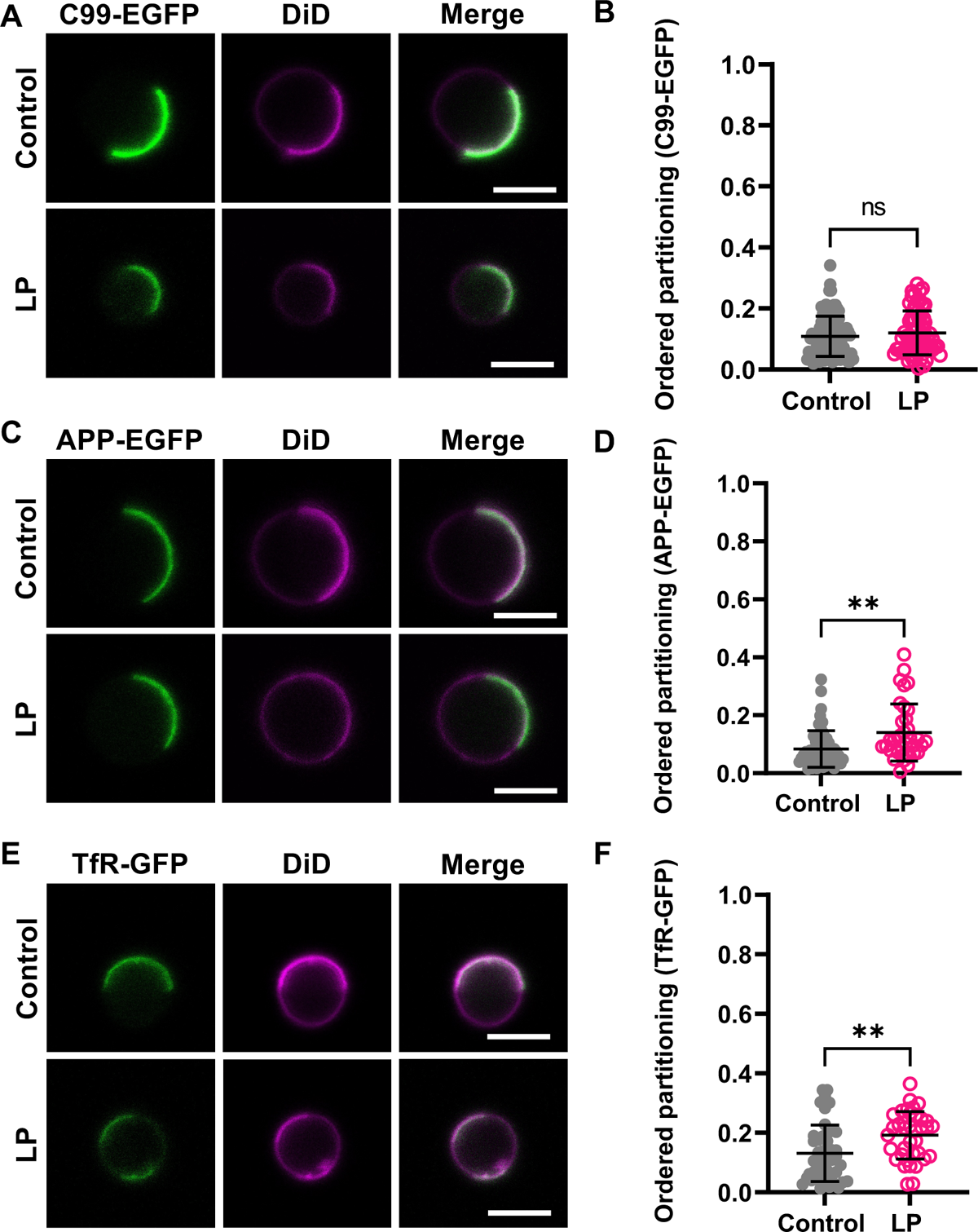
Non-raft proteins remain associated with disordered domains following lipid peroxidation. (**A**) Representative images of C99-EGFP in HeLa cell GPMVs. (**B**) Quantification of ordered partitioning of C99-EGFP-GFP across multiple GPMVs. Each data point corresponds to an individual GPMVs. Data are presented as mean ± SD for 65-102 GPMVs. (**C**) Representative images of APP-EGFP in HeLa cell GPMVs. (**D**) Quantification of ordered partitioning of APP-GFP across multiple GPMVs. Data are presented as mean ± SD for 38-54 GPMVs. (**E**) Representative images of TfR-GFP in HeLa cell GPMVs. (**F**) Quantification of ordered partitioning of TfR-GFP across multiple GPMVs. Data are presented as mean ± SD for 35-46 GPMVs. **, P < 0.01; ****, P < 0.0001 for unpaired two-tailed t-test. All data are representative of 2 independent experiments. Scale bars, 5 μm.

## Discussion

In the current study, we used GPMVs as a model to investigate how lipid peroxidation driven by the Fenton reaction impacts the properties of lipid rafts in biological membranes. Our findings reveal that lipid peroxidation significantly impacts several aspects of raft homeostasis (**Fig. 6**).

**Figure 6.**
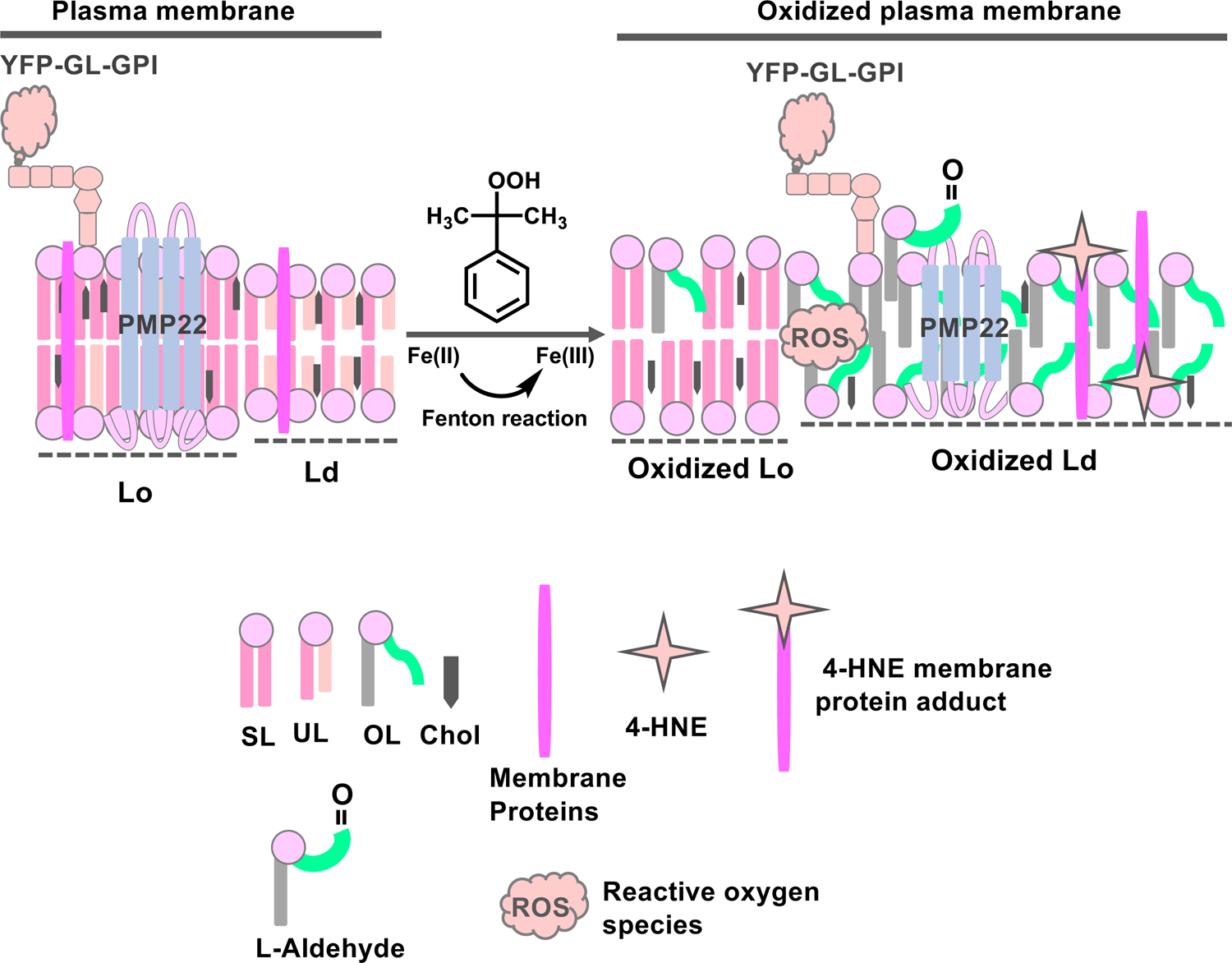
Working model for how lipid peroxidation induced via the Fenton reaction impacts raft and non-raft domains in biological membranes. Peroxidized lipid and their bioactive products such as 4-HNE preferentially accumulate in disordered domains. This is accompanied by increases in the relative abundance of the disordered phase, decreased lipid packing in both phases, and changes in protein composition in both phases as the result of selective redistribution of proteins from the ordered to the disordered phase. SL, saturated lipid, UL, unsaturated lipid, OL, oxidized lipid, chol, cholesterol.

One of the most striking effects of lipid peroxidation is the dramatic enhancement of membrane demixing into co-existing ordered and disordered domains, resulting in nearly all vesicles becoming phase separated. To our knowledge, this represents among the largest response to perturbation ever reported in the GPMV model. Several factors likely contribute to the heightened propensity of the membrane to phase separate. The most obvious is the accumulation of peroxidized lipids and their products in the disordered phase. This finding is in some ways not unexpected given that non-raft domains are generally thought to be enriched in lipids with unsaturated chains, including PUFAs. Furthermore, we observed a significant increased area fraction of the disordered phase and decreased lipid packing therein in peroxidized GPMVs. We also detected evidence for covalent modifications of membrane proteins and their segregation into non-raft domains by the bioactive aldehyde 4-HNE. The presence of this and other byproducts of lipid oxidation could also contribute to disruptions in membrane structure and decreased lipid packing.

Importantly, changes in membrane properties following lipid peroxidation were not limited to non-raft domains: peroxidized lipids and decreased lipid packing were also observed in raft phases. Although this result may seem surprising, it is consistent with reports that some PUFAs can be incorporated in rafts [64, 119]. Other lipid components of rafts may serve to dampen the progression of lipid peroxidation. For example, sphingomyelin has been reported to serve inhibit oxidative damage by preventing propagation of lipid peroxidation [120, 121], and enhanced levels of cholesterol have been reported to reduce membrane peroxidation in fibroblasts from patients with familial Alzheimer’s disease and to suppress lipid peroxidation in tumor cells [122, 123]. In future studies, it will be important to elucidate how the effects of lipid peroxidation depend on factors that influence membrane lipid composition such as growth conditions, cell cycle stage, nutritional sources, and disease state.

We also found that multiple proteins, including CTxB, GPI-anchored proteins, and multipass transmembrane proteins were dislodged from rafts upon peroxidation into non-raft domains. In contrast, non-raft proteins remained localized correctly despite the extensive accumulation of the peroxidized lipids in the non-raft phase. Thus, lipid peroxidation causes a loss of segregation of raft from non-raft proteins. The finding that raft proteins are selective mislocalized implies that their affinity for ordered phases is especially sensitive to changes in membrane structure that occur in response to lipid peroxidation. The exact mechanism underlying this sensitivity is not yet clear, but could be linked to loss of PUFAs and/or changes in the local structure of ordered raft domains, both of which have been suggested to aid in protein sorting [69, 70, 124]. Changes in protein structure induced by covalent modifications by bioactive products of peroxidation [21–23] could also contribute to the selective loss of protein affinity for the more ordered phase. Indirect mechanisms could also be involved [125]. How widespread the consequences of oxidation-dependent displacement of proteins from rafts are on cellular structure and function remains to be determined. This could be especially important for signaling pathways that are regulated by raft formation, raft targeting mechanisms, or segregation of raft and non-raft proteins [126, 127]. Another critical question is how the structure and function of individual membrane proteins are impacted by oxidized lipids. Evidence is already beginning to accumulate that oxidized lipids modulate the activity of transmembrane proteins whose function is sensitive to their membrane environment, such as G protein coupled receptors (GPCRs) [128]. Such effects could not only be important for proteins that normally prefer to function in a raft environment, but also for those that normally operate in non-raft domains.

Lipid peroxidation can be initiated by a variety of mechanisms [6]. In the current study, we used a non-enzymatic approach to generate reactive oxygen species driven by the Fenton reaction. The Fenton reaction occurs in biological systems and is thought to contribute to ferroptosis, a form of cell death triggered by iron-dependent lipid peroxidation [25, 26]. Importantly, recent evidence suggests that plasma membrane lipids are among those that undergo peroxidation during ferroptosis [129, 130]. Thus, disturbances in plasma membrane raft homeostasis are likely to be among the cellular defects that occur as ferroptosis is initiated and executed. PUFAs are also susceptible to react with reactive oxygen species which are produced by enzymatic (e.g., 12/15 lipoxygenase) processes, as well as those produced by photosensitizers and other mechanisms [23, 131]. An important goal for the future will be to establish whether these differing peroxidation mechanisms have similar or distinct effects on raft homeostasis.

In conclusion, our study illustrates that lipid peroxidation has profound effects on both raft and non-raft domains in biological membranes, including their stability, abundance, and lipid and protein composition. These findings suggest disruptions of membrane phase behavior may play a previously unrecognized role in ferroptosis and contribute to the pathology and progression of diseases linked to oxidative stress. Ultimately, understanding the consequences of lipid peroxidation on cellular membranes and their organization may offer new avenues to therapeutically target or exploit oxidative stress in various disease contexts.

## Materials and Methods

### Safety

No unexpected, new, and/or significant hazards or risks are associated with the reported work.

### Materials

DTT was purchased from Research Products International (Mount Prospect, USA) and was made fresh for every experiment. Paraformaldehyde was purchased as a solid from Fisher Scientific (catalog # AC416780250) and prepared as a 4% stock solution in H_2_O. All other chemicals such as CaCl_2_, HEPES, NaCl, Cumene hydroperoxide (catalog # 247502), and Iron (II) perchlorate hydrate 98% (catalog # 334081-5G) was purchased from Sigma. 18:0 NBD-PE (NBD-DSPE) 1,2-distearoyl-sn-glycero-3-phosphoethanolamine-N-(7-nitro-2-1,3-benzoxadiazol-4-yl) (ammonium salt) was purchased from Avanti Polar Lipids (catalog # 810141). DiD’ oil (catalog # D307), DiIC18(5) (catalog # D12730), BODIPY 581/591 C11 (catalog # D3861), and Di-4-ANEPPDHQ (Di4) (catalog # D36802) were purchased from Thermo Fisher Scientific. DiIC12 (1,1′-didodecyl-3,3,3′,3′-tetramethylindocarbocyanine perchlorate) (catalog # D383) and Fast DiI (catalog # D7756) were purchased from Invitrogen (Life Technologies). DAPT was obtained from Selleckchem (catalog # S2215). Cumene hydroperoxide and iron II perchlorate were each prepared as 10 mM stock solutions in H_2_O and stored at 4° C.

Antibodies used in this study include anti-myc Alexa Fluor 488 conjugate mAb (mouse) (Cell Signaling Technology, (catalog **#**2279S), rabbit anti-4 hydroxynonenal antibody (Abcam, catalog **#**ab46545), and Alexa Fluor 488 donkey anti-rabbit IgG (H+L) (Invitrogen, catalog **#**A21206). Cholera toxin subunit B (recombinant), Alexa Fluor 555 conjugate was from Themo Fisher Scientific (catalog **#**C22843). cDNA constructs used in this study include YFP-GL-GPI (model GPI-anchored protein; raft marker) [132]; TfR-GFP (transferrin receptor; non-raft marker), a gift from Dr. Ilya Levental [118]), and N41Q PMP22 (raft-associated multi pass transmembrane protein) [81]. C99-EGFP was as previously described [83] and was a gift from Dr Paola Pizzo [133]. APP-GFP was also a gift from Dr Paola Pizzo [133].

### Cell culture

WT-HeLa, COS-7, and RPE-1 cells were acquired from ATCC (Manassas, USA). Cells were cultured in Dulbecco’s modified Eagle medium (DMEM) containing 10% fetal bovine serum (FBS) and 1% pen/strep at 37 °C and 5% CO_2_. For all studies, cells were freshly plated a day prior to the preparation of GPMVs such that they would be 75-85% confluent on the day of the experiment.

### Transfection

For experiments using transfected cells, approximately 24 h prior to transfection, cells were plated to be 40–50% confluent at the time of transfection. Cells were transfected using FuGene-HD Transfection reagent (Promega, catalog #E2311) with a FuGene: DNA ratio of 3:1 using 4 μg of DNA in OptiMEM Reduced Serum Medium (ThermoFisher Scientific Catalog #11058021). The transfection medium was removed from cells ∼6–8 h post transfection, and cells were washed with Dulbecco’s modified Eagle medium and fresh culture media was added to each plate. GPMVs were prepared 24 h post transfection.

For experiments using C99-EGFP or APP-EGFP, twenty-four hours after transfection, cell culture media was exchanged for fresh media supplemented with 20 μM DAPT. DAPT was also included in all buffers used for GPMV preparation.

### Preparation and imaging of GPMVs

GPMVs were prepared as previously described [72]. In brief, cells were washed twice with GPMV buffer (2 mM CaCl_2_ /10 mM HEPES /0.15M NaCl, pH 7.4), followed by two washes with GPMV active buffer (GPMV buffer plus 25 mM formaldehyde and 2 mM DTT). Cells were then incubated with GPMV active buffer for 2 h at 37° C with shaking at 95-100 RPM. The GPMV-containing supernatant was decanted into a centrifuge tube and allowed to settle for 1 h at RT or overnight at 4° C. Aliquots of GPMVs (100 μL) were transferred from the bottom of the tube to a fresh tube for labeling or lipid peroxidation. We observed no difference in GPMV quality whether we imaged immediately or 24 h post-GPMV preparation.

To perform lipid peroxidation experiments, 100 μL of GPMVs were treated with a combination of 500 μM cumene hydroperoxide and 50 μM iron (II) perchlorate to serve as a source of ROS. GPMVs were post labelled with 1 μg/ml DiD (Ld phase marker [92, 93], diluted from a 1mg/mL stock in ethanol) to mark this disordered phase and 0.5 μg/ml NBD-DSPE (Lo phase marker, diluted from a 1mg/mL stock in ethanol) to mark the ordered phase. 30 minutes prior to imaging, 100 μL of GPMV solution was sandwiched between two coverslips coated with bovine serum albumin (BSA: 1mg/mL solution stock solution).

For some experiments, GPMVs were prepared from cells subjected to lipid peroxidation. Here, cells were incubated in the presence of 500 μM cumene hydroperoxide and 50 μM iron (II) perchlorate for 30 min and washed prior to GPMV preparation. A subset of GPMVs were then further incubated with 500 μM cumene hydroperoxide and 50 μM iron (II) perchlorate prior to labeling and imaging as described above.

GPMVs were imaged at RT using a Zeiss LSM 880 confocal microscope using a 63X oil objective. For most experiments, the confocal pinhole was set to 63.1 nm for all the channel and gain set to 700 for three different channels. Fluorophores were excited using the 488 nm line of an argon laser (NBD-DSPE), the 540 nm line of a HeNe laser (BODIPY 581/591) and the 633 nm line of a HeNe laser (DiD). Images were collected at a 1X digital zoom with a 1024 × 1024 or 512 × 512-pixel resolution for all the measurements and at 8–10X digital zoom for quantifying phase partitioning with a 1024 × 1024 or 512 × 512-pixel resolution.

### Detection of lipid peroxidation

The lipid peroxidation sensor C11-BODIPY (581/591) (Invitrogen, catalog # D3861) used to detect lipid peroxidation. For these experiments, GPMVs were first subjected to lipid peroxidation with 500 μM cumene hydroperoxide and 50 μM iron (II) perchlorate at RT or left untreated. They were then labeled with 1 μM of C11-BODIPY (581/591) and allowed to settle for 15-20 min prior to imaging at RT by confocal microscopy. For some experiments, GPMVs were labeled with C11-BODIPY (581/591), immediately followed by DiD.

### Di4 fluorescence lifetime imaging

GPMVs were prepared from HeLa cells as described above. They were then either left untreated or incubated with 500 μM cumene hydroperoxide and 50 μM iron (II) perchlorate. Di4 was added to a final concentration of 1 μg/ml from a stock solution of 1 mg/mL Di4 in ethanol. The GPMVs were then sandwiched between two coverslips coated with 1 mg/mL BSA solution and allowed to incubate for an additional 20 min prior to lifetime measurements.

Di4 lifetime images were performed on a Leica TCS SP8 system consisting of a Leica DMi8 inverted scanning confocal microscope with time-correlated single photon counting (TC-SPC) system and an 80 MHz-pulsed supercontinuum white light laser which allows continuously tunable excitation in the visible range coupled with filter-free detection. Samples were imaged using an HC PL APO CS2 63X/1.20 water objective. Di4 was excited at 488 nm and the emission was collected in the 550-800 nm range. The photon count rate was kept under 0.5 photons per laser pulse by adjusting the laser power, and sufficient frames were cumulatively acquired to obtain at least 100 photons per pixel. The fluorescence decay curves were fit with a bi-exponential re-convolution function adjusted to the instrument response function, and the average intensity-weighted lifetime was calculated for individual vesicles or area of interest and represented in the FLIM images as Di4 lifetime. Data is reported from two independent experiments. Statistical analyses were performed using GraphPad Prism 9.3.0.

### Immunofluorescence labeling of 4-HNE and PMP22

For 4-HNE labeling, GPMVs were prepared from HeLa cells as described above. Anti-4 hydroxynonenal antibody (Abcam, catalog # ab46545) was added to the GPMVs solution at a 1:500 dilution and gently agitated in the dark at RT for 1-2 h. GPMVs were subsequently labelled with a 1:500 dilution of Alexa Fluor 488 Donkey anti-Rabbit IgG (H+L) (Invitrogen, catalog # A21206) and gently agitated in the dark at RT for a minimum of 2-3 h. The GPMV preparation was split in half. Half of the labeled GPMVs were left untreated. The other half were incubated with a combination of 500 μM cumene hydroperoxide and 50 μM iron (II) perchlorate for 30 minutes. Finally, DiD was added to a final concentration of 1 μg/ml from a stock solution of 1 mg/mL in ethanol. The labeled GPMVs were then sandwiched between two coverslips coated with 1mg/mL BSA separated by a 1 mm silicon spacer and allowed to settle for 30 min at RT prior to imaging. As controls, either the primary antibody or secondary antibody was excluded from the labeling reactions.

For PMP22 labeling, GPMVs derived from HeLa cells transiently expressing PMP22 were then decanted into a 6 well plate. Alexa 488 mouse anti-myc mAb (Cell Signaling Technology, catalog # 2279S) was added to the solution at a 1:500 dilution and gently agitated in the dark at RT for at 3-5 h. GPMVs were then collected, transferred to a fresh tube, and allowed to settle for 1-2 h at RT. Next, GPMVs were either left untreated or subjected to lipid peroxidation as described above, then labelled at RT with 0.5 μg/ml Fast DiI (diluted from a 0.5 mg/mL stock in ethanol) to mark the disordered phase. GPMVs were mounted for imaging as described above. Imaging of GPMVs containing PMP22 was performed 4-6 h after labeling.

### CTxB labeling

GPMVs were prepared from COS-7 cells and either left untreated or subjected to lipid peroxidation using the procedures described above for HeLa cells GPMVs. Alexa Fluor 555-cholera toxin subunit B (Invitrogen, catalog # C22843) was added to 100 µl of GPMVs at a 1:100 dilution of a 1 mg/ml stock. The GPMVs were post labelled with 0.5 μg/ml NBD-DSPE (diluted from a 1mg/mL stock in ethanol) and 1 μg/ml DiD (diluted from a 1 mg/mL stock in ethanol) to mark the ordered and disordered phases, respectively, mounted, and imaged as described above.

### Quantification of Phase Partitioning

P_ordered_ was calculated as previously described [72, 82, 106] as Eq. 1

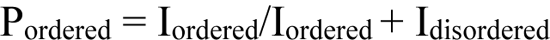

where I_ordered_ and I_disordered_ are the fluorescence intensity of a given fluorescent dye or protein in the ordered and disordered phases, respectively. To determine the phase of each domain, GPMVs were labeled with either an ordered membrane phase marker (NBD-DSPE), a disordered membrane phase marker (DiD), or both. ImageJ was used to manually perform a line scan across individual GPMVs to determine the highest fluorescent intensity at each pixel. The line scans were smoothed using a moving average (5 to 10 pixels) in Excel then the peak values were used to calculate P_ordered_ values. A P_ordered_ > 0.5 indicates a preference for the more ordered phase, and P_ordered_ < 0.5 corresponds to a disordered phase preference.

### Calculation of percentage of phase separated vesicles

Where indicated, the percentage of phase separated vesicles was calculated from tiled images using a semi-automated Matlab-based program, VesA [82]. Data from the green channel was used for the analysis.

### Statistical analysis

GraphPad Prism 9 was used to carry out statistical analyses.

## Author Contributions

The manuscript was written through contributions of both authors. Both authors have given approval to the final version of the manuscript.

## ACKNOWLEDGMENTS

We thank Dr. Paola Pizzo for providing constructs, members of Dr. Ilya Levental and Dr. Kandice Levental’s laboratory for assistance with the Di4 measurements, Yelena Peskova for technical support and assistance with image analysis, and Dr. Ajit Tiwari and Dr. Ilya Levental for feedback on the manuscript. This work was supported by NIH 1RF1 AG056147. The content is solely the responsibility of the authors and does not necessarily represent the official views of the National Institutes of Health.

## ABBREVIATIONS

4-HNE: 4-hydroxynonenal

APP: amyloid precursor protein

CTxB: cholera toxin B subunit

DiD1,1’: dioctadecyl-3,3,3’

3’: tetramethylindodicarbocyanine perchlorate

Di4: Di-4-ANEPPDHQ

FLIM: fluorescence lifetime imaging microscopy

GPMV: giant plasma membrane vesicle

I_disordered,_: fluorescence intensity of a given fluorescent dye or protein in the disordered phase

I_ordered,_: fluorescence intensity of a given fluorescent dye or protein in the ordered phase

Ld: liquid disordered

Lo: liquid ordered

NBD-DSPE: 1,2-distearoyl-sn-glycero-3-phosphoethanolamine-N-(7-nitro-2-1,3-benzoxadiazol-4-yl

PMP22: Peripheral Myelin Protein 22

P_ordered,_: ordered phase partitioning fraction

PUFA: polyunsaturated fatty acid

TfR: transferrin receptor

## ASSOCIATED CONTENT

### Supporting Information

Supporting information includes six supplementary figures showing both Fe (II) and cumene hydroperoxide are required for lipid peroxidation to occur in GPMVs; representative control experiments for 4-HNE staining; an example of how fluorescence intensity was quantified to calculate P_ordered_; lipid peroxidation enhances phase separation in RPE1-derived GPMVs; and lipid peroxidation causes the translocation of YFP-GL-GPI from ordered to disordered domains.

