## Supplementary Figures for "Lipid peroxidation drives liquid-liquid phase separation and disrupts raft protein partitioning in biological membranes"

**Figure S1.**

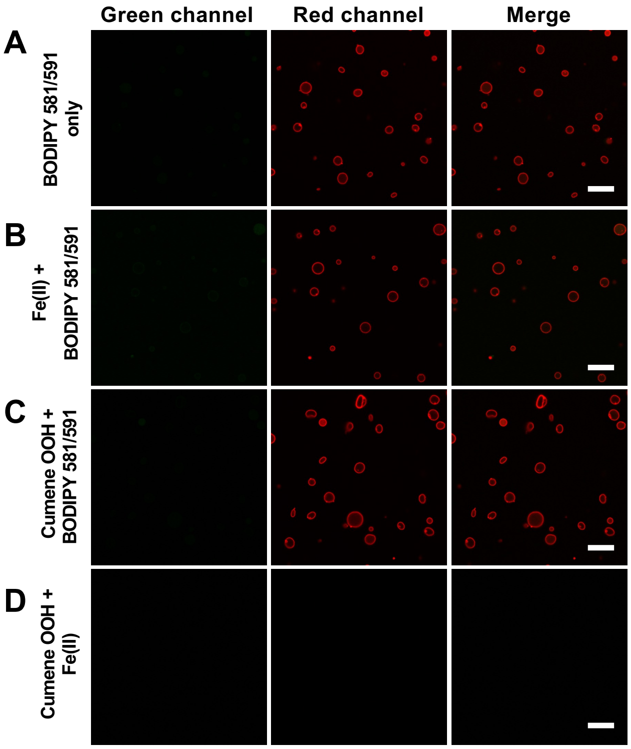


**Figure S1. Both Fe (II) and cumene hydroperoxide are required for lipid peroxidation to occur in GPMVs.** HeLa cell-derived GPMVs were incubated with the following reagents prior to imaging: (**A**) BODIPY 581/591 (1 µM); (**B**) Fe(II) (50 µM) followed by BODIPY 581/591 (1 µM); (**C**) Cumene hydroperoxide (500 µM) followed by BODIPY 581/591 (1 µM); and (**D**) Fe(II) (50 µM) and cumene hydroperoxide (500 µM) in the absence of BODIPY 581/591. No fluorescence was observed in the green channel when one or both lipid peroxidation reagents were omitted from the reaction, and no fluorescence was detected in either the green or red channels in the absence of BODIPY 581/591. Scale bar, 20 µm.

#### Figure S2.

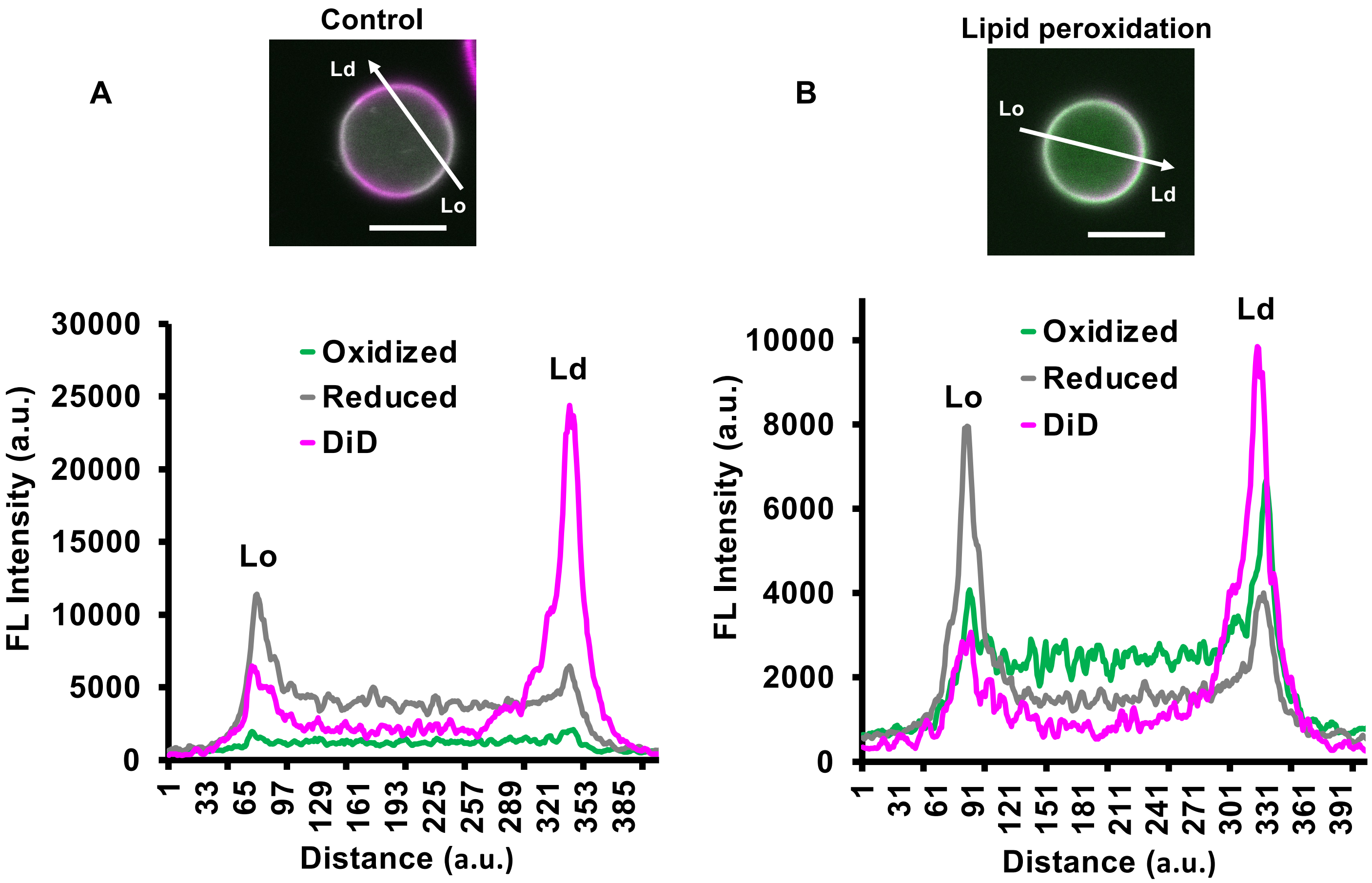


**Figure S2. Example of how fluorescence intensity was quantified to calculate the ratio of oxidized and reduced forms of BODIPY 581/591 in each phase.** Representative examples are shown for (**A**) a control GPMV and (**B**) a GPMV subjected to lipid peroxidation. A line bisecting each GPMVs was manually positioned to pass through both the Ld domain (enriched in DiD) and Lo domain (depleted in DiD). Plots of the fluorescence intensity along the line for all three fluorescence channels was generated. The peaks of the line plots were used to quantify the ratio of oxidized and reduced forms of BODIPY 581/591 in each phase as described in the Materials and Methods. a.u., arbitrary units. Scale bars, 5 µm.

### Figure S3.

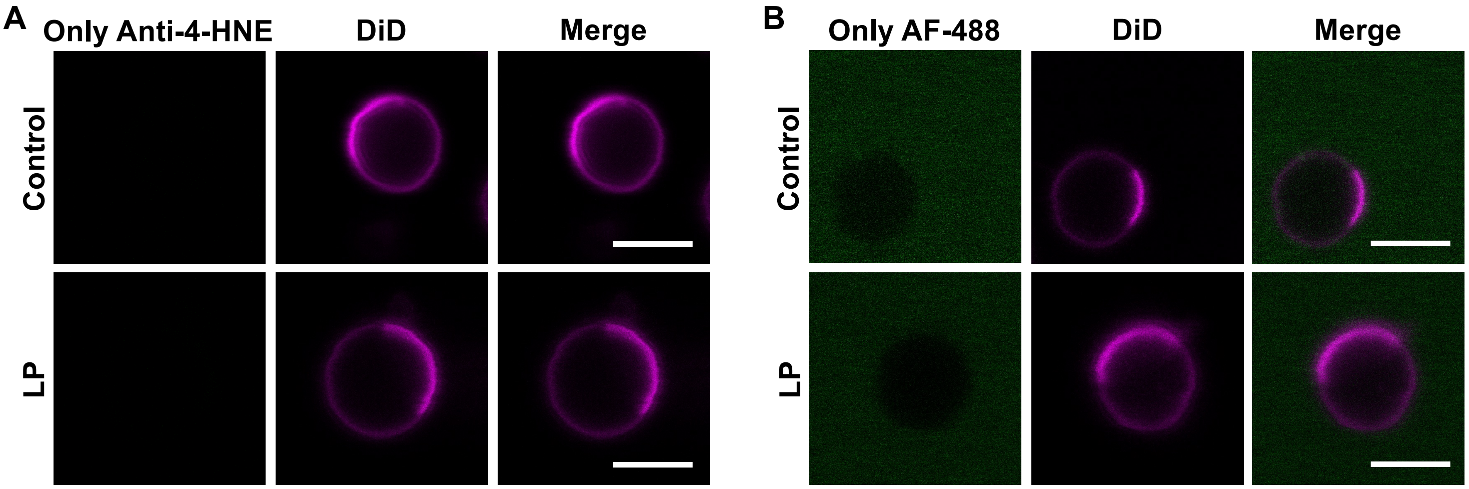


**Figure S3. Representative control experiments for 4-HNE staining.** GPMVs were either left untreated or subjected to lipid peroxidation, immunolabeled with an anti-4-HNE antibody or Alexa-488 secondary antibody, and then stained using DiD. Examples of representative GPMVs are shown. **(A)** Control verifying lack of fluorescent antibody staining in the absence of secondary antibody. **(B)** Control verifying lack of fluorescent antibody staining in the absence of primary antibody. Scale bars, 5 μm.

### Figure S4.

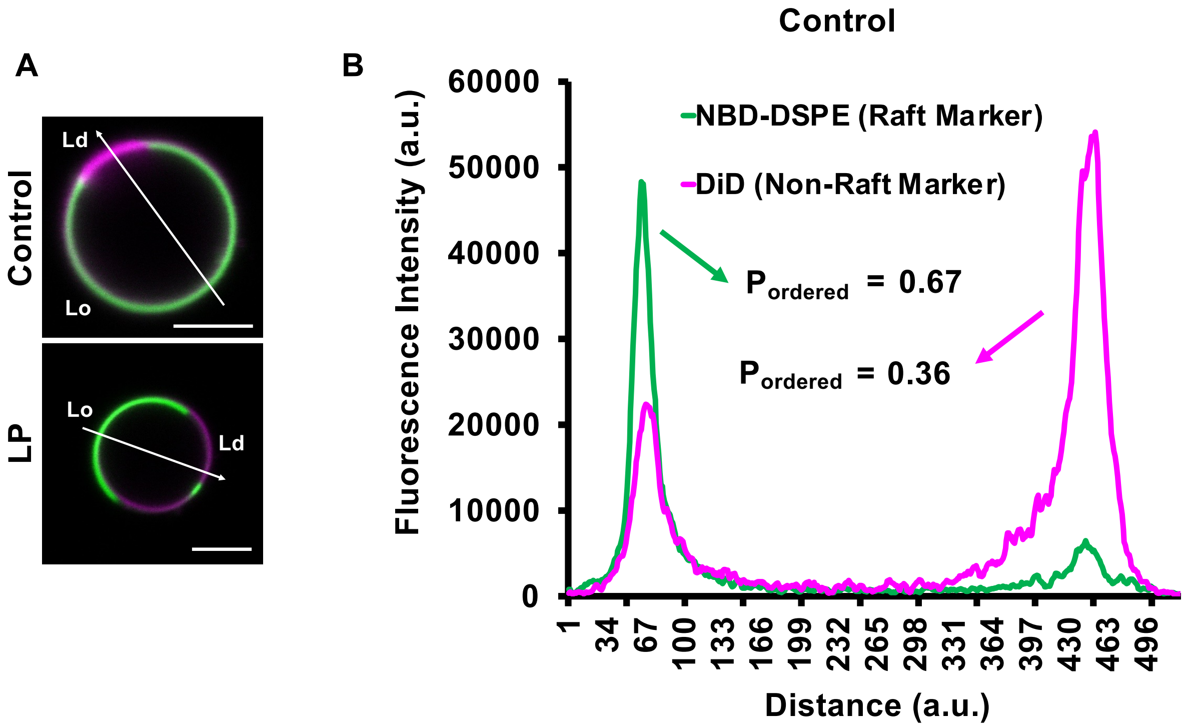


**C**

$$P_{ordered} = \frac{I_{ordered}}{I_{ordered} + I_{disordered}}$$

**D**

**$P_{ordered}$ > 0.5, Ordered phase preference**

**$P_{ordered}$ < 0.5, Disordered phase preference**

**$P_{ordered}$ = 0.5, No phase preference**

**Figure S4. Example of how fluorescence intensity was quantified to calculate $P_{ordered}$.** (**A**) Representative examples are shown for a control GPMV and a GPMV subjected to lipid peroxidation. A line bisecting each GPMVs was manually positioned to pass through both the Ld domain (enriched in DiD) and Lo domain (enriched in NBD-DSPE). Scale bars, 5 µm. (**B**) Plots of the fluorescence intensity along the line for both fluorescence channels was generated. The peaks of the line plots were used to quantify $I_{ordered}$ and $I_{disordered}$ for each probe. a.u., arbitrary units. (**C**) Equation used to calculate $P_{ordered.}$ (**D**) Definitions of $P_{ordered}$ values.

**Figure S5.**

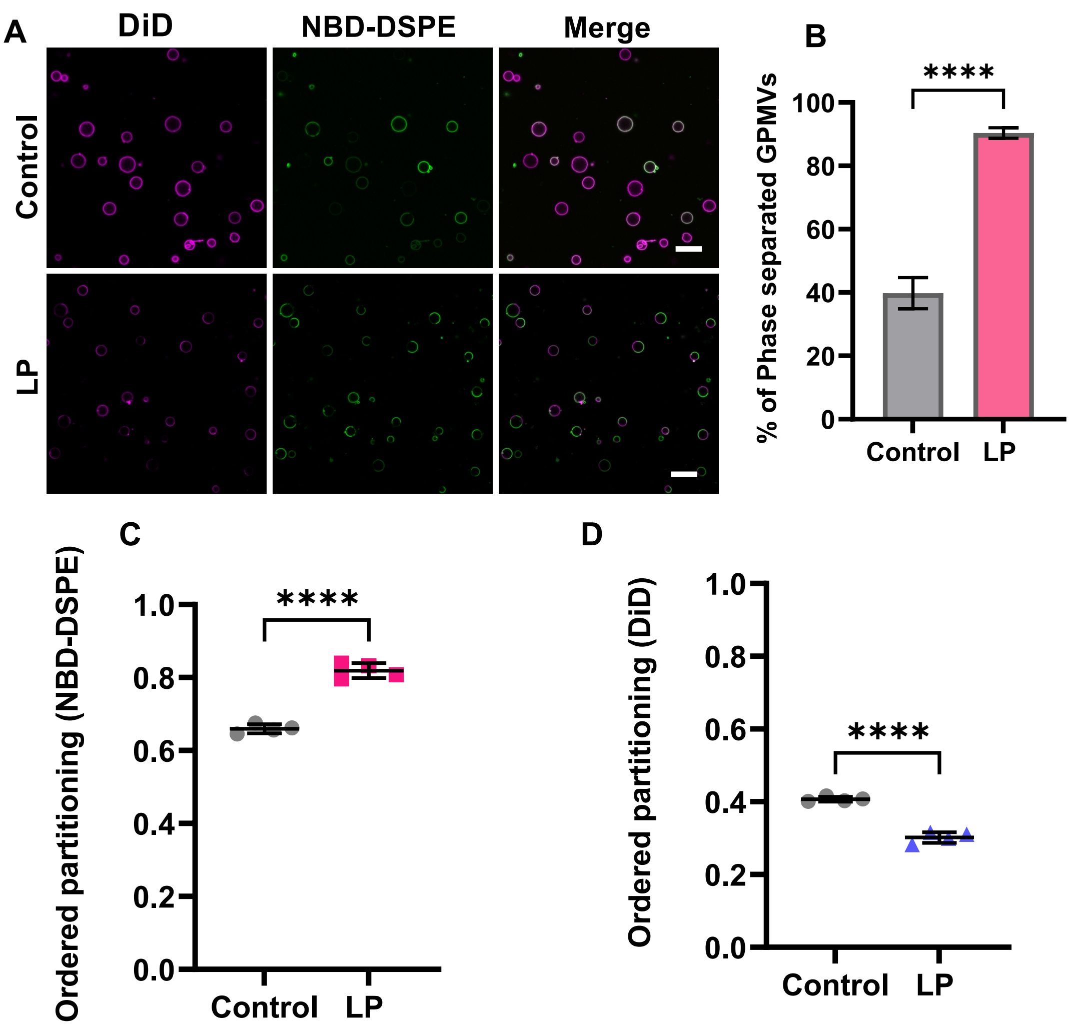


**Figure S5. Lipid peroxidation enhances phase separation in RPE1-derived GPMVs.** (**A**) GPMVs were either left untreated (control) or subjected to lipid peroxidation (LP). They were then labeled sequentially with NBD-DSPE (green) and DiD (magenta) prior to imaging at RT using confocal microscopy. Scale bar, 20 µm. (**B**) Quantification of the percentage of phase separated GPMVs for control versus lipid peroxidation conditions. The % of phase separated GPMVs was calculated using the green channel using VesA software. Bars correspond to mean ± SD for >100 GPMVs per condition. (**C, D**) Impact of lipid peroxidation on ordered partitioning of raft (NBD-DSPE) and non-raft (DiD) reporter dyes. Points in C and D represent >50 GPMVs in each group. ****, $P < 0.0001$ using unpaired two-tailed t-test.

### Figure S6

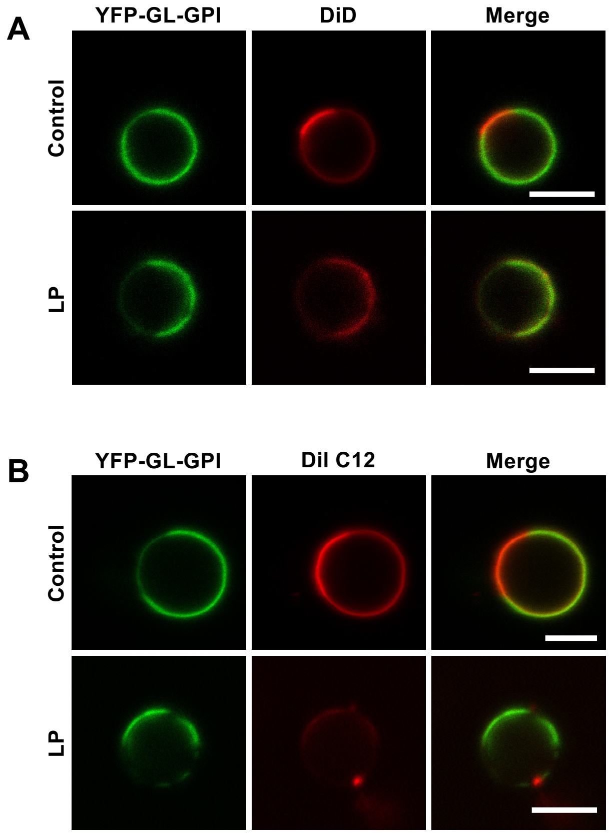


**Figure S6. Lipid peroxidation causes the translocation of YFP-GL-GPI from ordered to disordered domains.** Representative images of YFP-GL-GPI in HeLa cell-derived GPMVs under control conditions and following lipid peroxidation. GPMVs were labeled with (**A**) DiD or (**B**) DiIC12 after lipid peroxidation. Scale bars, 5 µm.
